# The Live Concert of Brains: Performer-Audience Neural Coupling Links Ensemble Coordination to Shared Audience Integration

**DOI:** 10.64898/2026.07.21.739936

**Authors:** Yue Ding, Jiameng Liu, Yifan Xu, Li Ji, Yubo Gao, Zheng Liang, Yingying Tang, Juan Huang, Xiaoqin Wang, Dan Zhang

## Abstract

Collective social events often require coordinated action by one group to become shared experience in another, yet how this transformation is organized across multiple brains remains unclear. Live music provides a tractable model of this problem because ensemble coordination, performer-audience alignment, and shared audience integration unfold within the same event. Here we tested whether performer-audience neural coupling acts as a cross-role neural interface linking ensemble coordination to shared audience integration. We recorded brain activity from a nine-person live performance system, consisting of a fixed three-performer ensemble and four independent audience groups, using synchronized multi-device functional near-infrared spectroscopy hyperscanning across live trio performance sessions. Inter-brain neural coupling was analyzed across three relational layers: performer-performer (PP), performer-audience (PA), and audience-audience (AA) coupling. Behavioral ratings showed strong affective engagement and shared evaluative alignment. Neural coupling during live performance was not expressed as a diffuse increase across channel pairs, but was organized into task-sensitive relational components with interpretable PC-corr network modules. Crucially, path-based mediation analyses revealed that PA components statistically bridged PP coordination and AA coupling, yielding multiple complete and partial PP → PA → AA pathways. Brain-behavior analyses further suggested that mediation-related PA components were linked to shared performance evaluation and emotional alignment. These findings identify performer-audience coupling as a cross-role neural interface through which coordinated production becomes linked to shared collective reception.

**Significant Statement:** How coordinated actions become shared experiences is a major problem in social neuroscience, yet most studies examine pairs of people. Using live music as a model of group interaction, we found that brain-to-brain alignment was organized across social roles rather than arising as a uniform response to the same event. Performer-audience alignment occupied a bridging position between coordination within the ensemble and integration within the audience. This identifies a systems-level architecture through which collective action may become linked to collective experience. The framework moves multi-brain research beyond dyads and offers a general approach for studying classrooms, public speaking, theater, rituals, and team events, where one group generates structured behavior that another group jointly receives and interprets.

## Introduction

Many natural social events require coordinated action by one group to be transformed into shared experience in another^1,2^. Public speaking, teaching^3^, theater, ritual^4^, sport^5^, and live music all involve structured behavior generated by agents in one social role and collectively received, interpreted, and evaluated by agents in another. Understanding such events requires moving beyond the question of whether two brains synchronize, and instead asking how inter-brain coupling is organized across differentiated social roles^6–9^. Live ensemble music provides a tractable model system for this question as coordinated production on stage and shared reception in the audience unfold within the same continuous event^10–12^. In live performance, performers coordinate timing, expression, and group-level musical structure in real time^13–16^, while audience members follow, evaluate, and affectively respond to the unfolding performance^17–19^. This makes live music a natural case of multi-level social coupling, in which coordinated production must be linked, through the performer-audience interface, to shared collective reception.

Such role-differentiated social events are unlikely to be captured by a single synchrony measure or by isolated dyadic interactions. They are organized by higher-order relational structures in which coordination, cross-role transmission, and collective alignment unfold across multiple interacting levels^6–8^. In live ensemble music, these levels can be mapped onto three forms of inter-brain coupling. Performer-performer (PP) coupling reflects coordination within the production system: musicians must align timing, predict co-performers’ actions, monitor errors, and adapt their own output to the ensemble^14–16,20^. Audience-audience (AA) coupling reflects convergence within the reception system, as listeners exposed to the same live event may align through shared attention, prediction, affective appraisal, and social context^13,21–23^. Performer-audience (PA) coupling occupies a distinct position between these two systems. It links agents who generate structured expressive action with agents who perceive, evaluate, and affectively respond to that action^24–27^. Thus, live music is not simply a unitary stimulus that drives synchrony in multiple brains, but a relational architecture in which performer coordination, performer-audience alignment, and shared audience integration may be functionally connected.

At the neural level, this relational architecture raises a central question: are PP, PA, and AA coupling independent consequences of the same acoustic event, or are they linked states within a single multi-brain system? Existing hyperscanning studies have provided important evidence for each component separately. Work on joint musical performance has shown coupling among performers during coordinated action^28–32^; studies of communication and performance reception have examined producer-receiver or performer-listener alignment^6,33,34^; and studies of collective listening have investigated neural convergence among listeners^21–23^. However, these lines of work have rarely been integrated within the same live event^35^. As a result, it remains unclear whether performer coordination on stage, performer-audience alignment across the stage-audience boundary, and shared audience integration are merely parallel responses to a common musical stimulus, or whether they form a structured cascade in which PA coupling links PP coordination to AA integration.

Performer-audience coupling is therefore a critical test case for moving from the presence of cross-role alignment to its systems-level function. Unlike coupling within a performing ensemble or within an audience, PA coupling spans a social boundary between agents who generate structured expressive action and agents who perceive, predict, evaluate, and affectively respond to that action. Prior studies have shown that producer-receiver or performer-listener neural coupling can accompany successful communication, social alignment, and music appreciation^33,34^. However, such findings have primarily established that cross-role neural alignment occurs and that it can be behaviorally meaningful. They do not determine whether PA coupling also functions as an intermediate relational state that connects coordination within a producer group to neural convergence within a receiver group. Live ensemble performance provides a tractable opportunity to test this possibility because production-side coordination, cross-role alignment, and reception-side convergence can be measured within the same unfolding social event.

Testing this bridging account requires a structured many-person design rather than a dyadic design. Dyadic hyperscanning can determine whether two individuals become neurally aligned, but it cannot reveal whether alignment across a producer-receiver boundary links coordination within a producer group to convergence within a receiver group^7–9,36^. A three-performer ensemble provides the minimal non-dyadic musical unit in which coordination among performers becomes a distributed group state rather than a single pairwise relation^16,37^. Pairing this fixed ensemble with independent audience groups further allows the same production-side coordination system to be observed across multiple reception contexts. This design therefore instantiates a tractable social topology with three relational layers: PP coupling as ensemble coordination, PA coupling as the cross-role interface, and AA coupling as shared audience integration. In this topology, the critical question is not whether more brains can be recorded simultaneously, but whether the PA interface occupies a mediating position between coordinated production and shared collective reception.

Testing this topology also requires an analytical framework that treats inter-brain coupling as a structured relational pattern rather than as a collection of isolated links. In a nine-person live performance system, each layer contains hundreds to thousands of channel-pair coupling features. A PA bridge between performer coordination and audience integration is therefore unlikely to be captured by any single channel pair; it should instead be expressed as a distributed coupling state linking the production and reception systems^8,9^. We therefore represented each relational layer as a low-dimensional component-level state. Principal component decomposition was used to identify task-sensitive coupling modes, and PC-corr was used to reconstruct the network modules underlying these modes by combining feature loadings with task-period co-fluctuation among coupling features^38^. This approach allowed us to preserve the relational architecture of the data while reducing its dimensionality, and to test whether live performance was organized as a PP → PA → AA cascade among component-level states rather than as independent pairwise synchrony effects.

Here we tested this proposal using synchronized multi-device functional near-infrared spectroscopy (fNIRS) hyperscanning during real live trio performances. A fixed three-performer ensemble was paired with four independent audience groups, allowing us to measure performer-performer coordination, performer-audience alignment, and audience-audience integration within the same live performance system. We quantified inter-brain coupling with wavelet transform coherence, represented task-sensitive coupling patterns as component-level relational states, and used path-based mediation analyses to test whether PA states statistically linked PP coordination to AA integration. This design allowed us to move beyond asking whether live music increases interpersonal synchrony, and instead ask whether coordinated production becomes linked to shared collective reception through a cross-role performer-audience neural interface.

## Results

### Live performance elicited shared affective and evaluative alignment

We first examined whether the live performance paradigm elicited subjective engagement and shared alignment across the performer-audience system. The experiment involved a fixed trio consisting of a bass guitarist, a keyboardist, and a clarinetist, paired with four independent audience groups (Fig. 1A left panel, Fig. S1A, Table S1). In each session, the trio performed three instrumental pieces spanning pop, jazz, and classical-crossover styles (Fig. 1A right panel, Table S2), while fNIRS signals were recorded simultaneously from performers and audience members. After each piece, performers and audience members independently rated performance quality and emotional engagement. To distinguish the intensity of subjective experience from its sharedness, we derived mean-rating indices, reflecting the common level of evaluation or engagement, and dyadic agreement indices, calculated as the negative absolute difference between two raters’ scores, with values closer to zero indicating greater convergence.

**Figure 1.**
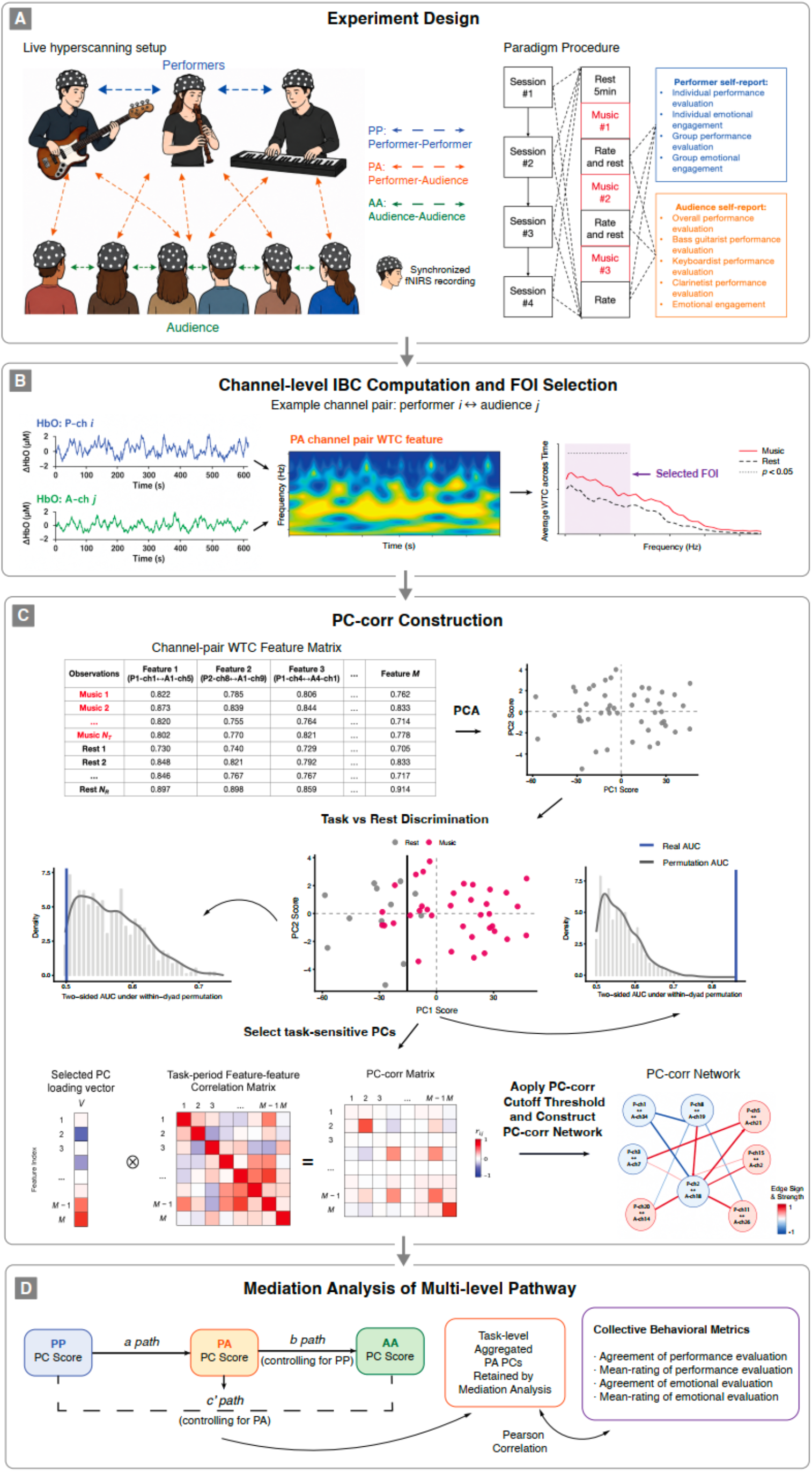
Experimental design and analytical framework. A, Experiment design. Upper panel: Live hyperscanning design; Middle panel: three relational levels including performer-performer coupling (PP), performer-audience coupling (PA), and audience-audience coupling (AA); Bottom panel: experiment procedure. **B, Channel-level coupling computation and frequency-of-interest selection.** For each channel pair, HbO time series were used to compute wavelet transform coherence (WTC), yielding a time-frequency representation of neural coupling. Live-performance-related coupling enhancement was quantified as the difference between live performance and rest. Frequencies showing reliable task-related enhancement were selected as frequencies of interest (FOIs) separately for PP, PA, and AA coupling. **C, PCA-based decomposition and PC-corr network construction.** FOI-averaged WTC values were organized into channel-pair feature matrices containing both task and rest observations. PCA was applied separately to PP, PA, and AA coupling matrices. Task-sensitive PCs were identified by their ability to discriminate live performance from rest, assessed against within-dyad permutation distributions of the AUC. For each retained PC, PC-corr networks were constructed by combining the PC loading vector with task-period feature-feature correlations, yielding interpretable network modules underlying task-sensitive coupling patterns. **D, Cascaded pathway and brain-behavior analysis.** Retained PP, PA, and AA PC scores were entered into a mediation framework testing whether performer-audience components mediated the pathway from performer coordination to shared audience coupling. The relationship of PA PC scores from complete mediation pathway and collective behavioral metrics was then explored.

The live performances were positively evaluated and emotionally engaging. Audience members reported high emotional engagement (M = 4.12, SD = 0.71) and high overall performance evaluation (M = 4.50, SD = 0.59), indicating strong subjective involvement. Mean-rating indices were also high across PP, PA, and AA layers, suggesting a generally favorable and emotionally engaging performance experience (Fig. S1B; Table S3). We next examined whether these ratings also converged across participants and relational layers. Among performers, agreement was stronger for ensemble-level ratings than for individual-level ratings. Agreement was highest for group performance evaluation and group emotional engagement (both M = −0.50), lower for individual performance evaluation (M = −0.72), and weakest for individual emotional engagement (M = −0.94). The weakest individual-level index, individual emotional engagement, was significantly lower than both group performance evaluation and group emotional engagement (t (35) = 3.16, p = 0.003, d = 0.53). Although the individual performance-evaluation contrast did not reach significance, it followed the same numerical pattern. Across the stage-audience boundary, performer-audience agreement was strongest for group emotional engagement (M = −0.90), exceeding agreement in performer-specific performance evaluation (M = −1.10, t (197) = −2.68, p = 0.008, d = −0.19) and overall performance evaluation (M = −1.05, t (197) = −2.36, p = 0.020, d = −0.17). Audience dyads also showed convergence in emotional engagement (M = −0.76, SD = 0.68) and overall performance evaluation (M = −0.61, SD = 0.57).

Together, these behavioral findings indicate that live performance generated not only favorable individual evaluations, but also shared affective and evaluative alignment across the performer-audience system. This provided the behavioral context for testing whether live performance was accompanied by structured neural coupling across PP, PA, and AA relational layers.

### Live performance organized inter-brain coupling into task-sensitive relational modules

We next asked how live performance changed neural coupling across the three relational layers. Inter-brain neural coupling was quantified using wavelet transform coherence (WTC) on HbO signals and computed separately for PP, PA and AA layers (Fig. 1B). To focus on task-relevant coupling, we first identified frequencies at which live performance showed stronger coupling than resting-state. Significant task-related coupling enhancement occurred primarily in low-frequency ranges below 0.2 Hz, yielding layer-specific frequencies of interest of 0.031 – 0.166 Hz for PP, 0.031 – 0.176 Hz for PA, and 0.020 – 0.130 Hz for AA (Fig. S1C). WTC values were then averaged within these FOIs, resulting in high-dimensional coupling matrices with 1,176 unique channel-pair features for PP, 2,304 features for PA, and 1,128 features for AA.

We then used the PC-corr framework^38^ to determine whether these high-dimensional coupling patterns were expressed as diffuse channel-pair changes or as organized relational modules. Within this framework, principal component decomposition served as the dimensionality-reduction step for extracting latent coupling modes from each layer, and each component was evaluated for its ability to discriminate live performance from resting state using a permutation-based AUC procedure (Fig. 2A). Task sensitivity was concentrated in a limited number of components rather than distributed across the full coupling space. For PP coupling, three principal components (PCs) survived the permutation threshold: PC1 (*AUC* = 0.928, *p* = 0.001), PC12 (*AUC* = 0.745, *p* = 0.014), and PC48 (*AUC* = 0.731, *p* = 0.013). PA coupling yielded 12 task-sensitive PCs, and AA coupling yielded 10 task-sensitive PCs; full AUC and permutation statistics are reported in Table S4.

**Figure 2.**
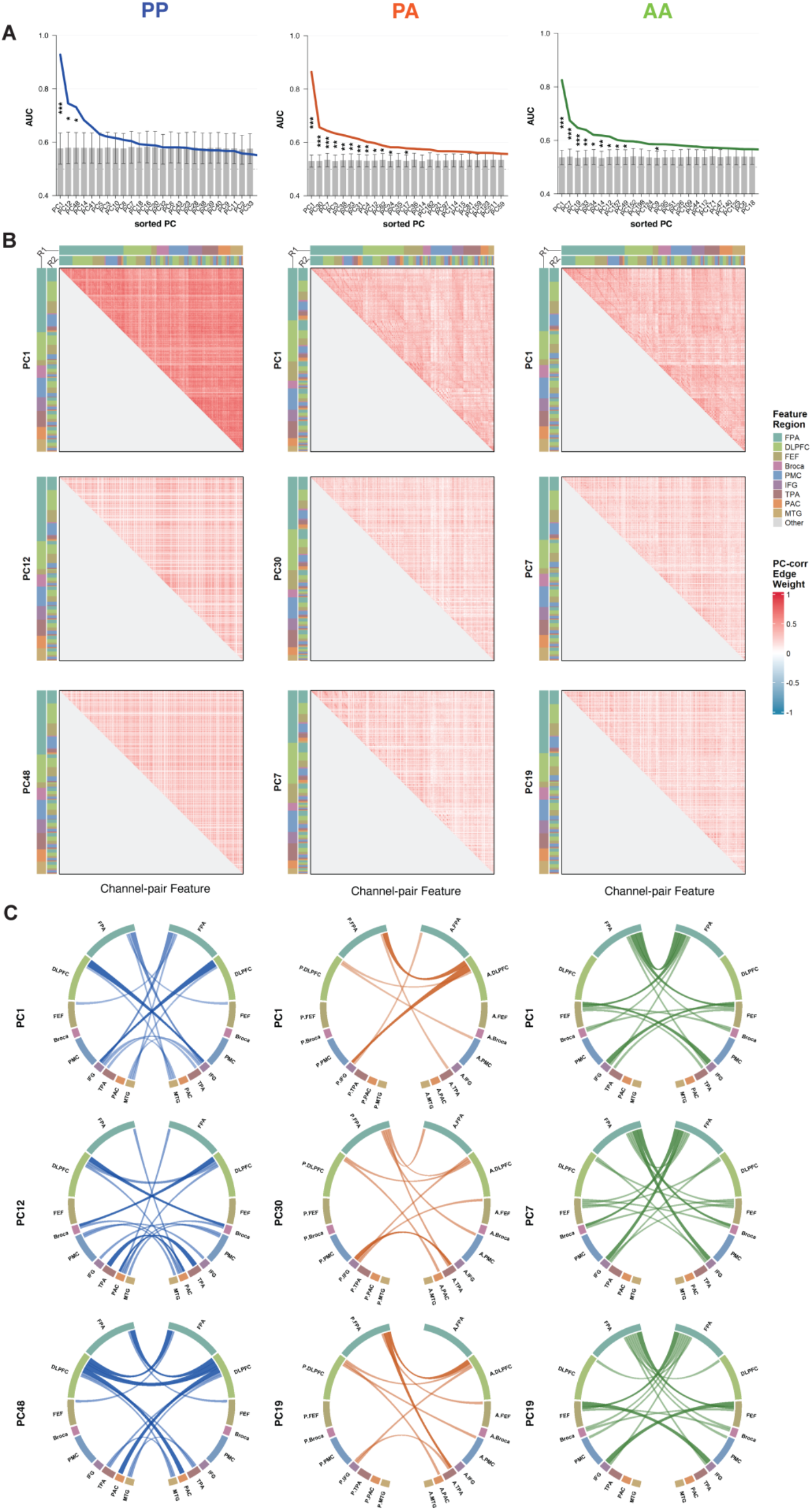
Live-performance-versus-rest discriminative PCs and corresponding PC-corr networks. A, Task sensitivity of PCs across the three relational layers. The three columns show PP, PA, and AA components, respectively. In each panel, the color line indicates the real AUC of each PC for discriminating live performance from rest, whereas the bars with standard deviations indicate the permutation-based AUC distribution. Asterisks indicate PCs that showed significant task sensitivity in the permutation test. *: *p* < 0.05, **: *p* < 0.01, ***: *p* < 0.001. **B, PC-corr matrices of the top three task-sensitive PCs.** For each coupling layer, the PC-corr matrices are shown for the three PCs with the highest task-discrimination AUCs. Each matrix represents PC-corr values by combining the PC loading of each channel-pair coupling feature with the task-period correlation between features. Each row and column in one matrix represents one network node, defined as a channel-pair coupling feature between two cortical regions. Each matrix element represents the PC-corr edge weight between two nodes, reflecting the extent to which two coupling features both contribute to the same task-sensitive PC and covary during live performance. Warmer colors indicate positive PC-corr weights, cooler colors indicate negative PC-corr weights, and color intensity reflects edge strength. Marginal color bars indicate the cortical regions associated with the channel-pair features. **C, Spatial distribution of the strongest PC-corr network nodes.** Circle plots show the channel-pair coupling features contained in the top 10 strongest PC-corr edges for the same PCs shown in B. Each circle represents a dyadic brain system, with the left and right halves corresponding to the two brains forming the coupling pair. For PA coupling, the left half represents the performer brain and the right half represents the audience brain. Lines indicate the cortical channel-pair coupling features that form the strongest PC-corr network nodes, thereby visualizing the spatial organization of the most prominent task-sensitive coupling modules.

For each retained task-sensitive component, PC-corr networks were reconstructed by combining the contribution of each channel-pair feature to the component with the task-period co-fluctuation among features. The resulting PC-corr heatmaps revealed organized blocks of feature co-fluctuation across PP, PA, and AA layers (Figure 2B; Fig. S2), indicating that live-performance-related coupling was supported by coordinated sets of coupling features rather than isolated channel pairs. The spatial distribution of the strongest PC-corr relations further showed layer-specific motifs involving prefrontal, frontal-motor, and temporal channel-pair features (Fig. 2C).

Together, these results indicate that live-performance-related neural coupling was not expressed as a uniform increase across all possible channel pairs. Instead, it was organized into a limited set of task-sensitive PC-corr modules, providing the component-level relational states used to test whether PA coupling statistically bridged PP coordination and AA integration.

### Performer-audience modules statistically bridged ensemble coordination and audience integration

We next tested whether PA coupling modules occupied a mediating position between performer coordination and audience integration. Retained task-sensitive PC-corr modules from the three relational layers were entered into a PP → PA → AA path-based mediation framework. Across all retained components, 360 PP × PA × AA combinations were tested. A path-based screen identified eight candidate cascades in which both the PP → PA path and the PA → AA path survived FDR correction.

Six cascades showed complete mediation patterns, whereas two showed partial mediation patterns (Fig. 3A and B). The complete mediation pathways involved three PP components, four PA components, and five AA components. PA PC1 appeared in two complete mediation pathways and linked both the dominant PP component (PP PC1 → PA PC1 → AA PC1: a = 0.584, b = 0.390, c’ = 0.089) and PP PC48 (PP PC48 → PA PC1 → AA PC1: a = 0.507, b = 0.462, c’ =-0.024) to the dominant AA component. Additional PA modules, including PA PC23 (PP PC1 → PA PC23 → AA PC7: a = 0.266, b = 0.208, c’ = 0.074), PA PC31 (PP PC12 → PA PC31 → AA PC34: a = 0.232, b = 0.196, c’ = 0.057), and PA PC4 (PP PC12 → PA PC4 → AA PC33: a =-0.195, b =-0.220, c’ =-0.054; PP PC12 → PA PC4 → AA PC14: a =-0.195, b = - 0.183, c’ =-0.125), mediated distinct pathways from performer coordination components to secondary audience-coupling components. In these complete mediation chains, the PP component predicted the PA component, the PA component predicted the AA component, and the residual direct PP → AA path was no longer significant after accounting for the PA bridge. The two partial mediation pathways (PP PC12 → PA PC12 → AA PC19: a = 0.325, b = 0.226, c’ = 0.178; PP PC1 → PA PC3 → AA PC1: a = 0.375, b =-0.171, c’ = 0.375) retained a significant direct PP → AA association, indicating that some PP-AA associations were not fully accounted by the intervening PA module.

**Figure 3.**
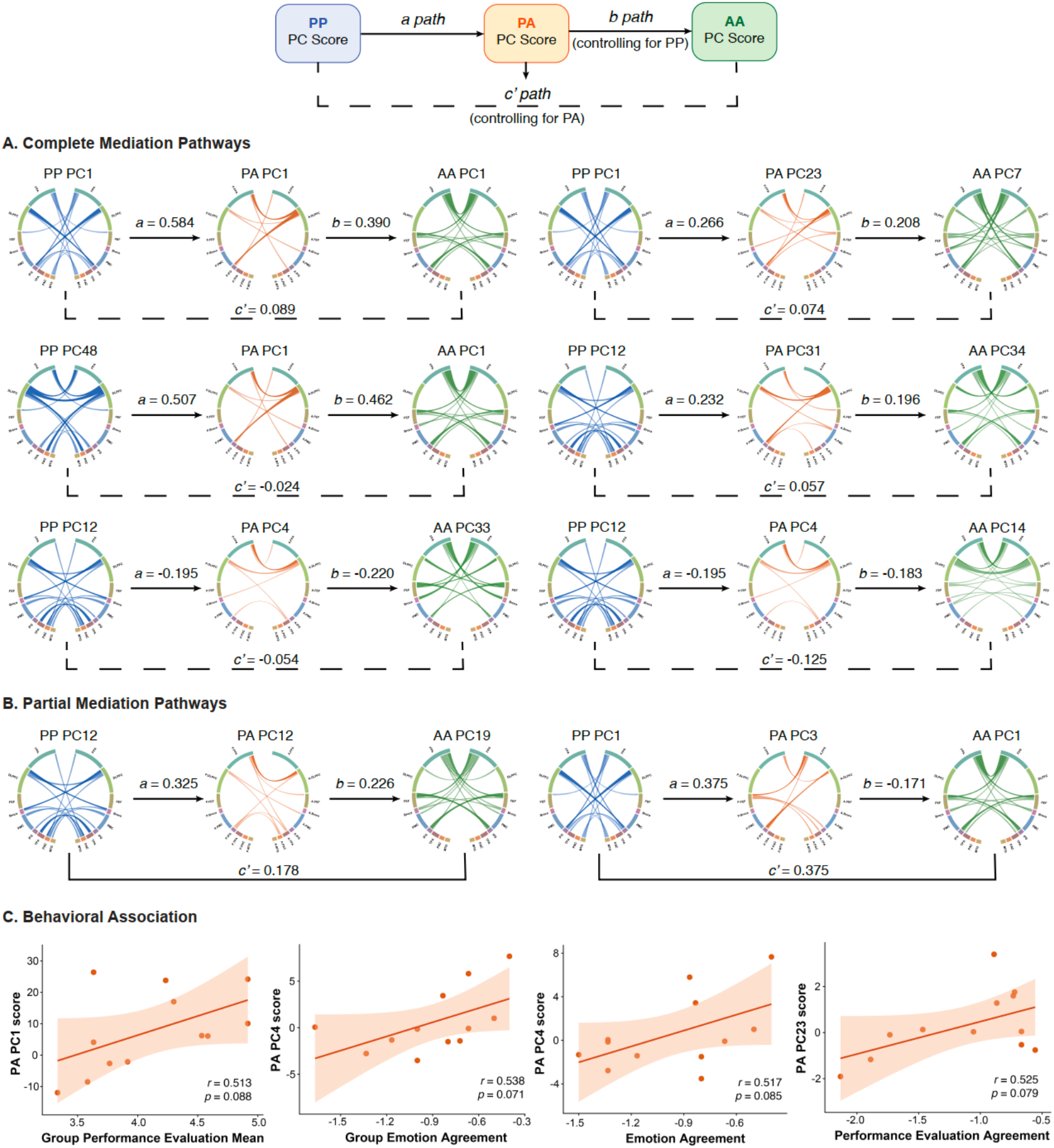
Performer-audience PC-corr modules mediate the cascade from performer coordination to shared audience coupling. The schematic at the top shows the mediation framework used to test whether performer-audience coupling modules statistically bridged performer-performer coupling and audience-audience coupling. The a path denotes the association between performer-performer and performer-audience PC scores, the b path denotes the association between performer-audience and audience-audience PC scores while controlling for performer-performer coupling, and the c′ path denotes the residual direct association between performer-performer and audience-audience PC scores after controlling for performer-audience coupling. **A, Complete mediation pathways.** Six PP → PA → AA cascades showed significant a and b paths after FDR correction, with a nonsignificant residual c′ path. Each pathway is shown with the corresponding PC-corr circle plots for the PP, PA, and AA modules. Arrows indicate the direction of the tested mediation paths, and path coefficients are shown next to each arrow. Dashed brackets indicate the residual direct c′ path. **B, Partial mediation pathways.** Two PP → PA → AA cascades showed significant a and b paths while retaining a significant residual c′ path, indicating that the PA module partially accounted for the relationship between performer coordination and shared audience coupling. **C, Behavioral associations of mediation-related PA modules.** Scatterplots show correlations between PA PC scores involved in significant mediation pathways and collective behavioral measures aggregated at the session-by-piece level. Behavioral indices include performer-audience mean rating of group performance evaluation, agreement in group emotional engagement, agreement in individual emotional engagement, and agreement in individual performance-evaluation ratings. Shaded areas indicate 95% confidence intervals. Agreement indices reflect dyadic alignment or convergence, whereas mean-rating indices reflect the common level of evaluation within a dyad.

The PC-corr circle plots embedded in the mediation diagram further clarified the architecture of these cascades. Performer-side modules were not reducible to a single coordination pattern: PP PC1, PP PC12, and PP PC48 each contributed to different mediation chains, suggesting multiple complementary modes of ensemble coordination. The PA modules did not simply duplicate either PP or AA networks. Instead, they formed distinct cross-role coupling configurations through which performer-side coordination was statistically associated with audience-side integration. At the audience layer, mediation pathways converged on both the dominant AA PC1 and several secondary AA components, indicating that shared audience coupling also emerged through multiple network modes rather than a single global synchrony pattern.

Together, these results support a PA-centered cascade in which performer coordination was statistically linked to audience integration through distinct performer-audience bridge modules.

### Brain-behavior analyses linked PA bridge modules to collective evaluation and affective alignment

Finally, we examined whether the PA modules implicated in the neural cascade showed preliminary behavioral relevance to the collective performance experience. Neural PC scores and behavioral measures were aggregated to the session-by-piece level, yielding 12 observations per analysis. This aggregation focused the analysis on task-level variation shared across the live performance system rather than on individual dyad-level noise.

Several mediation-related PA components showed convergent marginal associations with collective behavioral evaluations (Fig. 3C). PA PC1 was positively associated with the performer-audience mean-rating index for group performance evaluation (*r* = 0.513, *p* = 0.088). PA PC4 was positively associated with agreement in group emotional engagement (*r* = 0.538, *p* = 0.071) and with agreement in individual emotional engagement (*r* = 0.517, *p* = 0.085). PA PC23 was positively associated with the agreement in performer-specific performance evaluation (*r* = 0.525, *p* = 0.079).

Although these exploratory associations should be interpreted cautiously, they provide convergent evidence that PA components implicated in the neural cascade were related to shared aspects of performance experience. Specifically, the PA modules that statistically bridged PP coordination and AA integration also tended to covary with collective performance evaluation and affective alignment across the performer-audience system.

## Discussion

Live music is often described as a shared experience, but how this sharing is organized across multiple interacting brains remains poorly understood^11,13^. By simultaneously recording neural activity from a fixed performer trio and independent audience groups during real live performances, we show that live music organizes inter-brain coupling as a layered relational system. Live performance elicited shared affective and evaluative alignment, and neural coupling was structured into task-sensitive PC-corr modules across performer-performer, performer-audience, and audience-audience layers. The central finding was that performer-audience modules statistically bridged performer coordination and audience integration, yielding a PA-centered PP → PA → AA cascade. These findings suggest that live performance does not merely synchronize multiple brains in parallel; it organizes neural coupling across social roles, with performer-audience coupling serving as a cross-role interface linking coordinated production to shared collective reception.

The key conceptual implication is that performer-audience coupling should be distinguished from coupling within a single social role. Coupling among performers reflected coordination within the production system: musicians had to align timing, anticipate co-performers’ actions, monitor the evolving musical structure, and adapt their own output to the ensemble^15,16,29,31^. Coupling among audience members reflected integration within the reception system: listeners did not coordinate overt actions with one another, but they were exposed to the same unfolding event and could converge through shared attention, prediction, affective appraisal, and social context^13,21–23^. PA coupling occupied a different relational position. It linked agents who had asymmetric roles, different task demands, and complementary functions within the live event. In this sense, PA coupling is especially informative because it connects the neural dynamics of action generation with those of collective reception^3,25,26,33,34^. The present findings therefore extend prior work on performer-performer coordination, producer-receiver alignment, and listener-listener convergence by placing these relational forms within a single live performance architecture.

The mediation results further clarify the system-level function of this cross-role interface. Prior work has shown that producer-receiver and performer-listener coupling can accompany communication, shared understanding, and music appreciation^6,33,34^. The present findings extend this view by asking not only whether cross-role neural alignment occurs, but where it is positioned within a larger multi-brain architecture. In several complete mediation patterns, PP components predicted PA components, PA components predicted AA components, and the residual PP → AA association was no longer significant after accounting for the PA bridge. This pattern suggests that PA coupling was not merely an additional pairwise synchrony effect. Instead, mediation-related PA modules formed intermediate relational states through which performer coordination was statistically linked to shared audience integration.

This interpretation should be understood as cascade-compatible rather than as evidence for causal transmission. The present design does not show that neural activity flows from performers to audience members, nor that PA coupling is the only mechanism linking production and reception. Rather, it shows that the live performance system contained a structured PP → PA → AA association pattern in which PA components occupied a mediating position. The exploratory behavioral associations were consistent with this interpretation: mediation-related PA components tended to covary with shared performance evaluation and emotional agreement. Thus, the same performer-audience components that bridged neural coupling layers also showed preliminary relevance to collective appraisal and affective alignment.

The PA bridge was also instantiated by spatially organized PC-corr motifs rather than an abstract statistical variable alone. Across the mediation pathways, performer-side modules involved prefrontal, frontal-motor, and temporal coupling features^39,40^, consistent with the demands of ensemble performance, including timing alignment, action prediction, monitoring of musical structure, and adaptive coordination^28–32^. Audience-side modules involved recurrent fronto-frontal and temporal coupling features, consistent with shared reception supported by common attention, prediction, and appraisal^21–23^. Most importantly, the mediation-related PA modules formed distributed cross-role configurations linking performer frontal and temporal channels with audience prefrontal, inferior-frontal/premotor, and temporal channels^33,35^. These configurations are compatible with the proposed role of PA coupling as an interface through which coordinated action and expressive cues generated on stage are mapped onto listeners’ attentional, predictive, evaluative, and affective processes. Because a PC-corr node denotes a channel-pair coupling feature rather than a single cortical locus, these spatial patterns should not be interpreted as region-specific activations. Instead, they suggest that the PA-centered cascade was instantiated by distributed coupling motifs across prefrontal, frontal-motor, and temporal systems.

The present findings also illustrate the value of treating many-person hyperscanning data as relational states rather than as collections of isolated dyadic links^9,36^. In a live performance system, each relational layer contained hundreds to thousands of channel-pair coupling features. Analyzing each feature independently would obscure the higher-order organization of the system and greatly inflate descriptive complexity. The PC-corr framework allowed us to identify task-sensitive coupling modes and reconstruct the coordinated sets of coupling features underlying each mode. This component-level representation was especially important for testing the PA bridge, because the link between performer coordination and audience integration was unlikely to reside in a single channel pair. Instead, it emerged as a distributed performer-audience coupling state that could be positioned within a PP → PA → AA relational cascade.

Beyond live music, this framework may provide a model for studying how natural social events organize neural coupling across differentiated social roles. Many collective settings, including teaching^3^, public speaking, theater^41^, ritual^4^, team performance, and sport^5^, involve coordinated action generated by one group and collectively perceived, evaluated, and internalized by another. Such events cannot be fully characterized by asking whether two individuals synchronize. They require asking how coupling patterns are organized across production, cross-role interface, and reception layers. From this perspective, performer-audience coupling in live music provides a tractable case of a broader problem in social neuroscience: how neural dynamics coordinated within one social role become linked to shared experience in another.

Several limitations should be noted. First, the PP → PA → AA pathways reported here are statistical cascades rather than evidence of causal transmission. The present design does not demonstrate that neural activity flows from performers to audience members, nor that PA coupling is necessary for audience-level integration. It also cannot fully dissociate inter-brain coupling from shared stimulus-driven entrainment^42,43^, because all participants were embedded in the same live musical event. The task-versus-rest comparison and mediation framework show that coupling was structured across social roles, but future studies using live-versus-recorded controls, perturbation designs, audio-feature regression, or time-lagged modeling will be needed to test whether changes in the performer-audience interface causally alter downstream audience coupling and collective experience. Second, the present study involved a fixed trio, four performance sessions, and 22 valid audience members. This design increased ecological validity and preserved a stable production-side coordination system while allowing the same ensemble to be paired with independent audience groups. However, it also limits generalizability across ensembles, musical styles, audience configurations, and performance contexts. In addition, the brain-behavior associations should be interpreted as preliminary because neural and behavioral measures were aggregated to 12 session-by-piece observations. Larger studies with richer behavioral phenotyping, including continuous affect ratings, movement and gaze measures, autonomic responses, and indices of social connection, will be needed to determine which aspects of collective experience are carried by PA bridge components.

Together, these findings suggest that live performance organizes inter-brain coupling across differentiated social roles. Within this organization, performer-audience coupling serves as a cross-role neural interface that statistically links coordinated production to shared collective reception. Testing whether this PA-centered cascade reflects a causal mechanism, and whether similar cross-role interfaces organize other forms of collective social experience, will be an important next step for multi-brain social neuroscience.

## Methods

### Participants

Two types of participants were recruited: performers and audience members.

The performer group consisted of three fixed members of an amateur trio, including a bass guitarist (male, 24 years old), a keyboardist (male, 36 years old), and a clarinetist (female, 22 years old). All three performers had received more than five years of formal instrumental training and had been playing together for more than one year, with regular rehearsals at least once per week. A fixed ensemble was used to ensure stable collaborative rapport and to minimize variability that might arise from changing performer combinations.

Twenty-four audience members (age range = 18–32 years, *M* = 24.75, *SD* = 3.39) were recruited from a young adult population in Beijing, China. Inclusion criteria were normal hearing, right-handedness, no history of neurological or psychiatric disorders, and no prior exposure to live performances by the trio. Individuals with more than three cumulative years of formal instrumental or vocal training were excluded, as were those currently taking medications that might affect neural activity. Audience members were randomly assigned to one of four independent performance sessions, with six audience members in each session (Detailed demographic information in Supplementary Materials Table S1).

After data quality control, two audience participants were excluded, resulting in a final sample of three performers and 22 audience members. All participants provided written informed consent before the experiment. The study protocol was approved by the institutional ethics committee of the Department of Psychological and Cognitive Sciences, Tsinghua University.

### Stimuli

Three instrumental musical pieces from different genres were selected as the performance repertoire, including pop (*The Lost Days*, ∼250s), jazz (*Fly Me to the Moon*, ∼200s), and classical crossover (*The Rose*, ∼230s). The use of stylistically diverse music was intended to increase acoustic variability and thereby improve the sensitivity of subsequent statistical analyses. All pieces were purely instrumental, without lyrics, to avoid potential confounds related to linguistic or semantic processing.

### Procedure

The experiment was conducted across four independent performance sessions. Each session involved the same three performers and a different group of six audience members. The sessions took place in an acoustically treated multipurpose room of approximately 80 m². The performers were seated on a small stage at the front of the room, while audience members were seated in a line facing the performers approximately 3 m away.

Each session followed the same procedure. During the preparation phase (approximately 30 min), participants were fitted with fNIRS optode caps and signal quality was checked. A resting-state baseline was then recorded for 5 min, during which all participants were instructed to sit still, remain relaxed, and stay awake. This was followed by the live music phase (approximately 35 min), during which the trio performed the three musical pieces in sequence. Audience members were instructed to listen naturally without performing any explicit task. During the closing phase (approximately 10 min), the equipment was removed and participants received compensation.

### Behavioral Measurements

Following each musical piece, performers and audience members independently completed brief subjective rating questionnaires using 5-point Likert scales.

Performers rated four items encompassing both individual-and group-level dimensions: (1) individual performance evaluation (“How would you rate the overall quality of your own performance just now?”, 1 = very poor, 5 = very good); (2) individual emotional engagement (“How would you rate your level of emotional engagement during your own performance just now?”, 1 = not engaged at all, 5 = fully engaged); (3) group performance evaluation (“How would you rate the overall quality of the ensemble’s performance just now?”, 1 = very poor, 5 = very good); and (4) group emotional engagement (“How would you rate the ensemble’s level of emotional engagement during the performance just now?”, 1 = not engaged at all, 5 = fully engaged). Individual-level items were intended to capture each performer’s perception of their own performance, whereas group-level items reflected each performer’s perception of the ensemble’s collective state.

Audience members rated five items: (1) overall performance evaluation (“How would you rate the overall quality of the ensemble’s performance just now?”, 1 = very poor, 5 = very good); (2) bass guitarist performance evaluation (“How would you rate the overall quality of the bass guitarist’s performance just now?”, 1 = very poor, 5 = very good); (3) keyboardist performance evaluation (“How would you rate the overall quality of the keyboardist’s performance just now?”, 1 = very poor, 5 = very good); (4) clarinetist performance evaluation (“How would you rate the overall quality of the clarinetist’s performance just now?”, 1 = very poor, 5 = very good); and (5) emotional engagement (“How would you rate your level of emotional engagement while listening just now?”, 1 = not engaged at all, 5 = fully engaged). The three performer-specific performance evaluation items were designed to capture audience members’ differentiated appraisals of each performer’s contribution, whereas the overall performance evaluation item reflected their holistic evaluation of ensemble quality with an emphasis on perceived coordination. The emotional engagement item reflected the subjective intensity of their affective experience during listening.

### Performer-Performer (PP) behavioral indices

For each performer dyad and each piece, inter-rater agreement on each of the four rating dimensions was quantified as the negative absolute difference between the two performers’ scores:

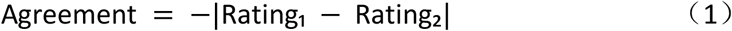

This transformation yields a value closer to zero as agreement increases, aligning directionally with neural synchrony indices (e.g., WTC) such that higher values reflect greater concordance. Four agreement indices were thus derived for each dyad per piece: individual emotional engagement agreement (*PP Emotion Agreement*), individual performance evaluation agreement (*PP Performance Evaluation Agreement*), group emotional engagement agreement (*PP Group Emotion Agreement*), and group performance evaluation agreement (*PP Group Performance Evaluation Agreement*). In addition, four mean indices were computed by averaging the two performers’ scores on each dimension: mean individual emotional engagement (*PP Emotion Mean*), mean individual performance evaluation (*PP Performance Evaluation Mean*), mean group emotional engagement (*PP Group Emotion Mean*), and mean group performance evaluation (*PP Group Performance Evaluation Mean*). With three performers forming three dyads across 12 pieces (three pieces × four sessions), this yielded 36 observations per agreement index.

### Performer-Audience (PA) behavioral indices

For each performer-audience dyad and each piece, agreement between a performer’s rating and an audience member’s rating was computed using the same formula:

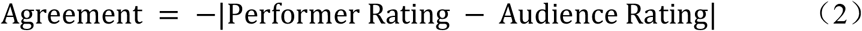

Here, the audience member’s performer-specific performance evaluation corresponding to the specific performer in the dyad (i.e., bass guitarist, keyboardist, or clarinetist performance evaluation rating) was used for *Individual Performance Evaluation Agreement*, whereas the audience member’s overall performance evaluation rating was used for *Group Performance Evaluation Agreement*. Four agreement indices were derived: individual performance evaluation agreement (*PA Individual Performance Evaluation Agreement*), group performance evaluation agreement (*PA Group Performance Evaluation Agreement*), individual emotional engagement agreement (*PA Individual Emotion Agreement*), and group emotional engagement agreement (*PA Group Emotion Agreement*). Correspondingly, four mean indices were computed by averaging the performer’s and audience member’s scores on each dimension: *PA Individual Performance Evaluation Mean*, *PA Group Performance Evaluation Mean*, *PA Individual Emotion Mean*, and *PA Group Emotion* Mean. With 22 audience members and 3 performers forming 69 dyads across 3 pieces per session, this yielded 198 observations per agreement index.

### Audience-Audience (AA) behavioral indices

For each audience dyad and each piece, inter-rater agreement on the rating dimensions was quantified as the negative absolute difference between the two audience members’ scores:

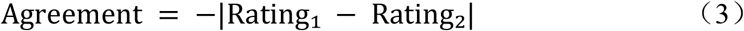

As with PP indices, two AA agreement indices were derived for each dyad per piece: emotional engagement agreement (*AA Emotion Agreement*) and overall performance-evaluation agreement (*AA Performance Evaluation Agreement*) In addition, two mean indices were computed by averaging the two audience members’ scores on each dimension: mean emotional engagement (*AA Emotion Mean*) and mean performance evaluation (*AA Performance Evaluation Mean*). With five to six audience members forming ten to fifteen dyads each session, this yielded 50 observations per agreement index.

### Data Recording

Neural activity was recorded using nine portable fNIRS systems (NirSmart, Danyang Huichuang Medical Equipment Co., Ltd.), with three devices assigned to the performers and six to the audience members. Each system used dual-wavelength near-infrared LED sources (760 and 850 nm) and photodetectors, and recorded at a sampling rate of 11 Hz with a source-detector separation of 3 cm. Each device provided 48 measurement channels covering bilateral prefrontal and temporal regions, which could be approximately mapped to the frontopolar area (FPA), dorsolateral prefrontal cortex (DLPFC), frontal eye field (FEF), premotor cortex (PMC), inferior frontal gyrus (IFG), superior temporal gyrus (STG), and middle temporal gyrus (MTG) on the basis of anatomical landmarks^44^.

Multi-device synchronization was achieved using a hardware-based triggering scheme. All nine fNIRS systems were connected in series to a common network switch, and a central control computer simultaneously broadcast UDP timestamps and trigger signals to all devices. Pre-experimental testing confirmed that the temporal synchronization error across devices was smaller than one sampling interval (< 91 ms). The audio playback system was synchronized with the fNIRS acquisition using the same trigger mechanism.

### Data Preprocessing

Raw fNIRS signals were preprocessed in Homer3^45^. The preprocessing pipeline consisted of five steps. First, channels with a signal-to-noise ratio (SNR) below 2 were excluded. Second, raw light intensity signals were converted into optical density. Third, motion artifacts were corrected using the SplineSG method^46^. Fourth, a band-pass filter of 0.01 – 0.5 Hz was applied. Finally, optical density was converted into oxygenated hemoglobin (HbO) concentration changes using the modified Beer-Lambert law^47^. Only HbO signals were retained for further analyses, as HbO is generally considered more sensitive to task-related cortical hemodynamic responses in fNIRS studies. To minimize edge artifacts, the first and last 15 s of each trial (165 sampling points at 11 Hz) were discarded prior to analysis.

## Data Analysis

### Neural coupling estimation

Interpersonal neural coupling was quantified using wavelet transform coherence (WTC)^48^. WTC is a time-frequency analysis approach that captures the phase consistency between two nonstationary signals across both time and frequency, making it particularly suitable for neural signals with dynamic temporal structure. It has been widely adopted in fNIRS hyperscanning research^9^.

For each channel, the HbO time series was subjected to continuous wavelet transform using the Morlet wavelet as the mother wavelet. The frequency range was set to 0.01 – 0.5 Hz to match the relevant frequency band of fNIRS signals. WTC was then calculated between two channels by smoothing the cross-wavelet spectrum and normalizing it by the product of the corresponding individual power spectra. WTC values ranged from 0 to 1, with higher values indicating stronger coupling at a given time-frequency point. To facilitate parametric statistical analysis, WTC values were transformed using Fisher’s Z transformation.

### Dyad organization

According to participant roles, all neural coupling measures were classified into three types.

Performer-performer coupling (PP) referred to inter-brain coupling between each pair of performers. Because there were three performers in each session, this yielded three performer dyads per session. The resulting coupling matrix was a symmetric 48 × 48 channel-pair matrix. Symmetric channel pairs (e.g., ch_i-ch_j and ch_j-ch_i) were averaged and mirrored to the complementary positions, resulting in 1,176 unique channel pairs.

Performer-audience coupling (PA) referred to inter-brain coupling between each performer and each audience member, yielding 3 × *n* audience dyads per session. This resulted in a full 48 × 48 non-symmetric matrix containing 2,304 channel pairs. Because performer and audience channels were not interchangeable, no symmetrization was applied.

For audience-audience coupling (AA), dyads were defined between pairs of audience members within the same session, following the same pairwise organization as PP coupling. Sessions with five valid audience members yielded 10 AA dyads, whereas sessions with six valid audience members yielded 15 AA dyads.

### Identification of frequencies of interest

For each coupling type, WTC values were computed separately for the task and resting-state conditions. Task-related frequencies of interest (FOIs) were identified in a data-driven manner. Across the full frequency range of 0.01 – 0.25 Hz, one-tailed paired *t* tests were then performed at each frequency point to test whether the task WTC was significantly greater than rest WTC across all channel pairs. These *p* values were corrected using the FDR method. Frequency ranges that survived correction in *t*-test analyses were defined as group-specific FOIs. This strategy avoided subjective a priori frequency selection while ensuring that the selected frequency ranges were robust for each coupling type.

### Principal component decomposition of task-related neural coupling patterns

Task-related neural coupling patterns were decomposed separately for PP, PA, and AA coupling. For each coupling level, WTC values were first averaged within the corresponding frequency range of interest for every dyad, session, condition, and channel pair. The resulting sample-by-feature matrix contained both task-period and resting-state observations, with rows representing dyad-level observations and columns representing channel-pair coupling features.

Before decomposition, channel-pair features with more than 10% missing values were removed, and the remaining missing cells were imputed using the median of the corresponding feature. The cleaned feature matrix was z-scored across observations and submitted to principal component analysis (PCA) using singular value decomposition. All available PCs were scanned rather than restricting the analysis to high-variance components.

Task sensitivity of each PC was quantified by its ability to discriminate task from rest observations. For each PC, a two-sided area under the receiver operating characteristic curve was computed as max(*AUC, 1-AUC*), thereby ignoring the arbitrary sign of the component. Statistical significance was assessed with 1,000 within-dyad permutations of the task/rest labels. PCs with permutation *p* <.05 were retained as task-sensitive components for downstream analyses. The sign of each retained PC was aligned such that the mean PC score during task periods was greater than the mean score during rest periods; therefore, higher aligned PC scores indicated more task-like coupling patterns.

After task-sensitive PCs were selected, PC-corr networks were constructed using the loading vector from the PCA fitted on the combined task/rest matrix, together with feature-feature correlations computed only from task-period observations. Thus, the resulting PC-corr network represents task-period co-fluctuation among channel-pair coupling features, constrained by their contribution to a PC that was task-sensitive relative to rest.

Following the official PC-corr implementation^38^, the loading vector *V* for each selected PC was first transformed as:

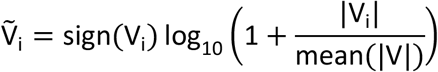

The transformed loadings were then rescaled to preserve sign while mapping their absolute values to the interval [0, 1]:

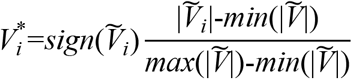

Features with |*V_i_*\*| greater than a predefined cutoff were retained. We examined cutoffs of 0.60, 0.65, and 0.70. For each retained pair of features *i* and *j*, the PC-corr edge weight was computed as:

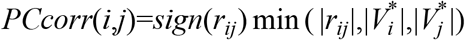

where *r_ij_* is the Pearson correlation between the two channel-pair features. Edges with |*PCcorr*(*i,j*)| below the cutoff were removed, and isolated nodes were excluded from the final network.

### Mediation analysis of multi-level neural coupling pathways

We next tested whether PA coupling components statistically mediated the relationship between performer coordination and shared audience neural coupling. The primary mediation chain was specified as PP → PA → AA. For each PA task observation, the score was taken directly from the corresponding performer-audience dyad. The PP score was computed as the mean of the PP dyads involving that performer, and the AA score was computed as the mean of the AA dyads involving that listener. This pairing yielded 198 task-period observations across four sessions, 12 performer identities nested within sessions, 22 audience members, and 66 PA dyads.

All retained PP, PA, and AA PCs were tested in a full factorial manner, resulting in 360 PP × PA × AA combinations. For each combination, variables were standardized and linear mixed-effects models were fitted to estimate the a path (PP → PA), the b path (PA → AA controlling for PP), and the direct path c’ (PP → AA controlling for PA). The default random-effects structure included random intercepts for performer and listener, with session included as a fixed effect. When this model was singular, a predefined fallback sequence with simpler random-effects structures was used.

Mediation candidates were defined using a path-based criterion: both the a path and the b path were required to survive FDR correction^49,50^. Among these candidates, mediation type was determined from the direct path c’. A candidate was classified as complete mediation when c’ was not significant after correction, and as partial mediation when c’ remained significant^51^.

### Brain-behavior association analysis

To examine whether task-sensitive coupling components were related to collective behavioral evaluations, neural PC scores and behavioral measures were matched at the appropriate dyadic level and then aggregated to the session-by-piece level. This yielded 12 observations per analysis cell, corresponding to four sessions and three musical pieces. The aggregation was used to focus on task-level variation shared across participants rather than trial-level dyadic noise.

The brain-behavior analysis focused on PA PCs retained by the path-based mediation screen. Pearson correlations were computed between aggregated PA PC scores and PA behavioral metrics, including subjective agreement and mean-rating indices for overall and emotional evaluations at both individual and group levels. Because this analysis was based on 12 aggregated session-by-piece cells, effects with *p* <.10 were treated as marginal associations.

## Data and Code Availability

The datasets and code used in the current study are openly available in OSF at https://osf.io/atcyg/overview?view_only=73601f100e9d4a3e867a360758643df4

## Acknowledgement

This work was supported by the National Natural Science Foundation of China (T2341003), and Shanghai Municipal Health Commission (2024ZZ2066). We thank Dr. Carlo V. Cannistraci for recommending the PC-corr algorithm. We thank all the participants.

## Author Contributions

D.Z. and X.W. conceptualized the study. J.L., Y.X. and L.J. were responsible for data collection. Y.D. and J.L. contributed to formal analysis and writing the original draft. D.Z. and Y.D. contributed to methodology, visualization, funding acquisition, and manuscript editing. Z.L. and Y.T. provided resources. All authors reviewed and approved the final manuscript.

## Ethical approval statement

The study was conducted in accordance with the Declaration of Helsinki and approved by the local ethics committee at the Department of Psychological and Cognitive Sciences, Tsinghua University (IRB No. THU202312). Participants gave their written informed consent and received monetary compensation for their participation. The publication of identifiable images of research participants was authorized with their informed consent.

## Artificial Intelligence (AI) Disclosure

Following the Generative AI Delegation Taxonomy (GAIDeT), the authors disclose the use of GPT-5.4 and Gemini-3.1 within the domain of Writing and Editing, specifically for proofreading and editing. The tool was used to improve grammar, wording, clarity, and readability of text originally prepared by the authors. It was not used for study conceptualization, literature review, research design, data collection or analysis, interpretation of results, or formulation of scientific conclusions. All AI-assisted revisions were critically reviewed and, where necessary, modified by the authors, who take full responsibility for the accuracy, integrity, and final content of the manuscript.

