## Supplementary material for "The Live Concert of Brains: Performer-Audience Neural Coupling Links Ensemble Coordination to Shared Audience Integration": SI Figure S1-2, Table S1-5

A

Representative photographs removed in this preprint version to protect participant privacy

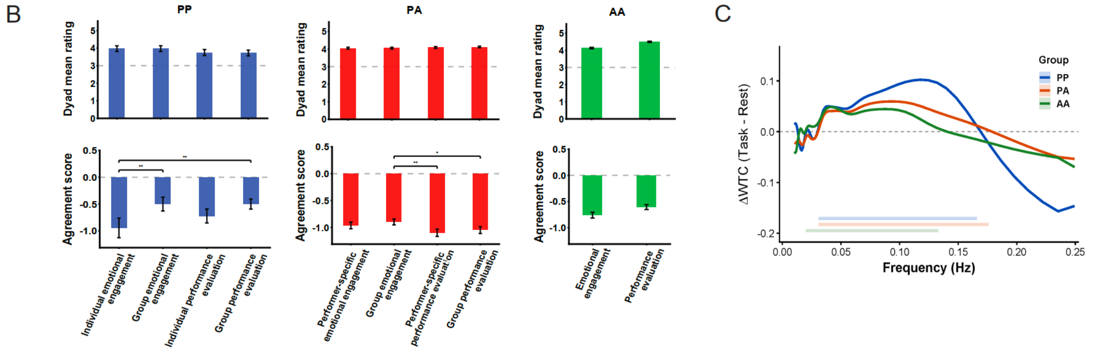


**Figure S1. A, Experimental setting of live fNIRS hyperscanning.** Representative photographs show the live trio performance setup (upper panel) and the audience arrangement (bottom panel). **B, Dyad mean and alignment of behavioral ratings during live performance.** Top panels show dyadic mean-rating indices, calculated as the average of the two raters' scores on the same behavioral dimension. Higher values indicate a stronger common level of emotional engagement or performance evaluation within the dyad. Bottom panels show dyadic agreement scores, calculated as the negative absolute difference between two raters' scores on the same behavioral dimension. Values closer to 0 indicate stronger agreement or convergence between raters. Behavioral indices are shown separately for performer–performer (PP), performer–audience (PA), and audience–audience (AA) dyads. PP includes individual and group emotional engagement, as well as individual and group performance evaluation. PA includes performer-specific and group emotional engagement, as well as performer-specific and group performance evaluation. AA includes emotional engagement and performance evaluation. Bars represent group means, and error bars indicate SEM. Dashed gray lines indicate the neutral midpoint of the 5-point rating scale for mean ratings and perfect agreement for agreement scores. PP, PA, and AA are shown in blue, red, and green, respectively. *$p < 0.05$, **$p < 0.01$. **C, Frequencies of interest for task-related neural coupling.** The curves show task-related coupling enhancement, defined as ΔWTC between task and rest, separately for performer–performer coupling (PP, in blue), performer–audience coupling (PA, in orange), and audience–audience coupling (AA, in green). Horizontal bars indicate the selected frequencies of interest (FOIs) for each coupling level, which were used for subsequent PCA, PC-corr network construction, and mediation analyses.

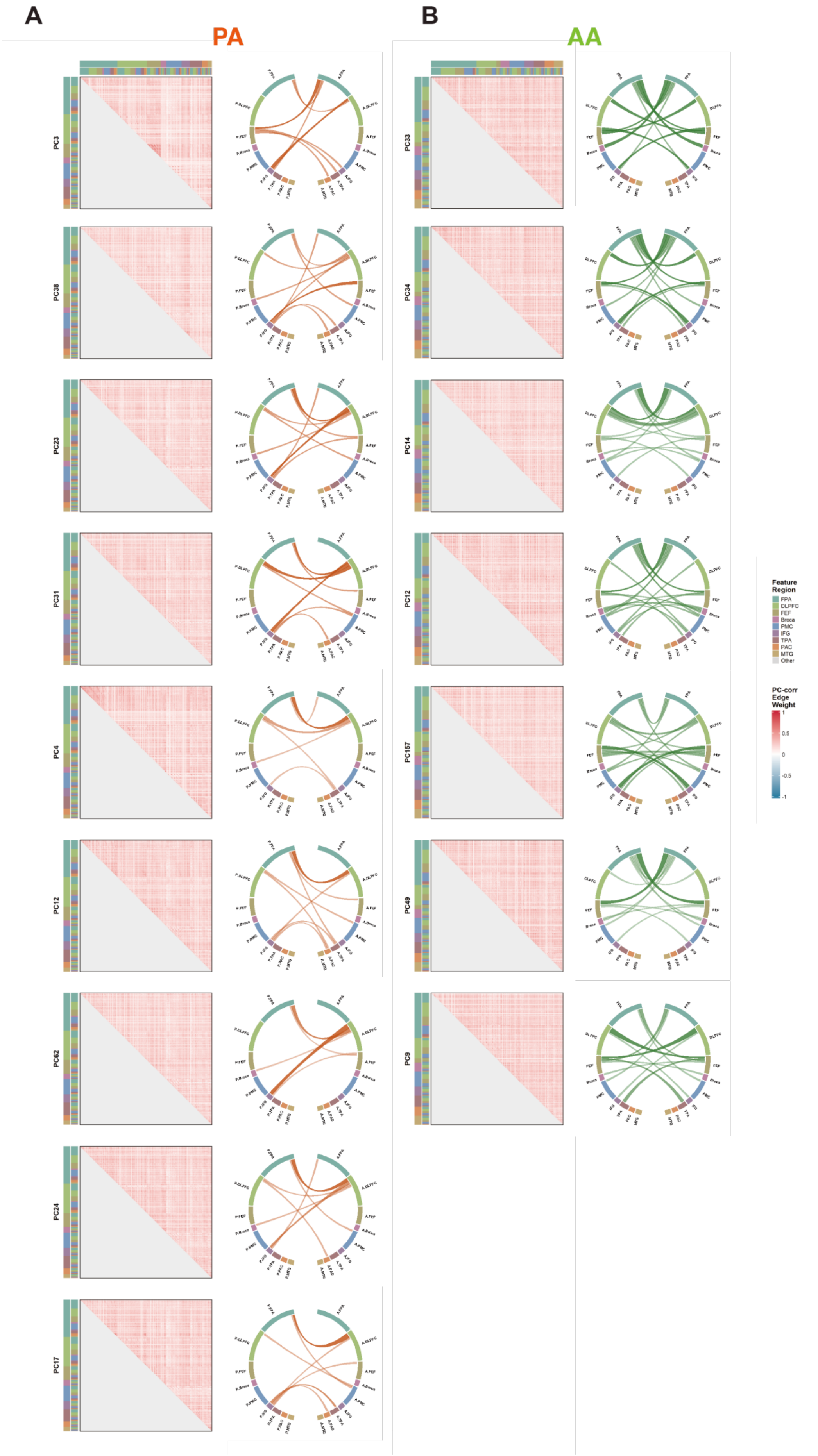
A
PA
B
AA
PC3
PC38
PC23
PC31
PC4
PC12
PC62
PC24
PC17
PC33
PC34
PC14
PC12
PC157
PC49
PC9
Feature Region
FPA
DLPFC
FEF
Broca
PMC
IFG
TPA
PAC
MTG
Other
PC-corr Edge Weight
1
0.5
0
-0.5
-1

**Figure S2. A, Remaining significant PCs in the PA group.** Each row shows one significant PC. The left panel shows the full PC-corr matrix, with channel-pair features ordered by brain-region assignment. The top and left color bars indicate the regions of the two channels composing each feature. Matrix color represents PC-corr edge weight, with red indicating positive weights, blue indicating negative weights, and gray indicating the masked lower triangle and diagonal. The right panel shows the chord diagram constructed from the top 10 nodes of the corresponding PC, ranked by matrix strength, defined as the row sum of absolute PC-corr edge weights in the full matrix. Orange links indicate PA connections. **B, Remaining significant PCs in the AA group.** The layout and annotations are the same as in A. Green links indicate AA connections. The shared legend denotes brain-region colors and PC-corr edge-weight values.

**Table S1. Demographic Characteristics of Audience Participants by Session**

| Session ID | *N* | Age (years) | Age *M(SD)* | Sex Ratio (M:F) |
|---|---|---|---|---|
| 1 | 6 | 24-29 | 25.50 (2.35) | 0:1 |
| 2 | 6 | 20-28 | 22.83 (3.13) | 1:1 |
| 3 | 6 | 18-31 | 25.00 (4.43) | 1:1 |
| 4 | 6 | 23-32 | 25.67 (3.44) | 2:1 |
| Total | 24 | 18-32 | 24.75 (3.39) | 5:7 |

**Table S2. Musical Repertoire and Retained Performance Duration**

| Name | Type | Duration mean (s) | Duration std (s) |
| --- | --- | --- | --- |
| The Rose | Classical crossover | 221.50 | 5.21 |
| The Lost Days | Pop | 243.75 | 5.55 |
| Fly Me to the Moon | Jazz | 183.43 | 5.71 |

**Table S3. Self-Report and Listener Rating Scores**

| Rating source | Rating dimension | *N* | *M (SD)* | Median | Range |
|---|---|---|---|---|---|
| Performer | Individual performance | 45 | 3.80 (1.04) | 4.00 | 1-5 |
| | Individual emotion | 45 | 3.98 (1.12) | 4.00 | 1-5 |
| | Group performance | 45 | 3.67 (0.98) | 4.00 | 2-5 |
| | Group emotion | 45 | 3.96 (1.02) | 4.00 | 2-5 |
| Listener | Overall performance | 66 | 4.50 (0.59) | 5.00 | 3-5 |
| | Emotion rating | 66 | 4.12 (0.71) | 4.00 | 3-5 |
| | Keyboardist performance | 66 | 4.62 (0.55) | 5.00 | 3-5 |
| | Bass performance | 66 | 4.05 (0.94) | 4.00 | 2-5 |
| | Clarinet performance | 66 | 4.56 (0.70) | 5.00 | 2-5 |

**Table S4. Significant AUC Results Based on Permutation Testing**

| Group | Principal component | Observed AUC | Permutation mean AUC | Permutation *p* value | Group | Principal component | Observed AUC | Permutation mean AUC | Permutation *p* value |
|---|---|---|---|---|---|---|---|---|---|
| **PP** | PC1 | 0.928 | 0.576 | .001 | PA | PC1 | 0.864 | 0.530 | .001 |
| | PC12 | 0.745 | 0.578 | .014 | | PC30 | 0.657 | 0.531 | .001 |
| | PC48 | 0.731 | 0.578 | .013 | | PC7 | 0.644 | 0.534 | .001 |
| **AA** | PC1 | 0.825 | 0.536 | .001 | | PC3 | 0.634 | 0.530 | .003 |
| | PC7 | 0.674 | 0.538 | .001 | | PC38 | 0.627 | 0.532 | .003 |
| | PC19 | 0.647 | 0.534 | .001 | | PC23 | 0.620 | 0.532 | .005 |
| | PC33 | 0.640 | 0.537 | .002 | | PC31 | 0.612 | 0.531 | .005 |
| | PC34 | 0.621 | 0.539 | .014 | | PC4 | 0.602 | 0.530 | .005 |
| | PC14 | 0.618 | 0.533 | .007 | | PC12 | 0.598 | 0.532 | .013 |
| | PC12 | 0.615 | 0.538 | .012 | | PC62 | 0.588 | 0.533 | .034 |
| | PC157 | 0.601 | 0.538 | .030 | | PC24 | 0.582 | 0.530 | .034 |
| | PC49 | 0.597 | 0.537 | .036 | | PC17 | 0.578 | 0.531 | .037 |
| | PC9 | 0.585 | 0.534 | .048 | / | | | | |

**Table S5. Correlations Between PA Principal Components and Behavioral Measures**

| PC | Behavioral measure | n | Pearson *r* | *p* value | PC | Behavioral measure | n | Pearson *r* | *p* value |
|---|---|---|---|---|---|---|---|---|---|
| PC1 | Group Mean | 12 | 0.513 | .088 | PC3 | Group Mean | 12 | 0.112 | .729 |
| | Group Emotion Mean | 12 | 0.446 | .147 | | Group Emotion Mean | 12 | 0.033 | .919 |
| | Emotion Mean | 12 | 0.425 | .169 | | Emotion Mean | 12 | 0.111 | .732 |
| | Individual Performance Evaluation Mean | 12 | 0.419 | .176 | | Individual Performance Evaluation Mean | 12 | -0.050 | .877 |
| | Group Performance Evaluation Agreement | 12 | 0.304 | .336 | | Group Performance Evaluation Agreement | 12 | 0.217 | .498 |
| | Emotion Agreement | 12 | 0.236 | .460 | | Emotion Agreement | 12 | -0.190 | .555 |
| | Group Emotion Agreement | 12 | 0.219 | .494 | | Group Emotion Agreement | 12 | -0.231 | .469 |
| | Individual Performance Evaluation Agreement | 12 | 0.165 | .608 | | Individual Performance Evaluation Agreement | 12 | 0.023 | .943 |
| PC4 | Group Mean | 12 | 0.106 | .742 | PC12 | Group Mean | 12 | -0.203 | .527 |
| | Group Emotion Mean | 12 | 0.010 | .974 | | Group Emotion Mean | 12 | -0.490 | .106 |
| | Emotion Mean | 12 | -0.055 | .866 | | Emotion Mean | 12 | -0.484 | .111 |
| | Individual Performance Evaluation Mean | 12 | 0.060 | .854 | | Individual Performance Evaluation Mean | 12 | -0.228 | .477 |
| | Group Performance | 12 | 0.182 | .572 | | Group Performance | 12 | -0.179 | .578 |

| | | | | | | | | | |
|---|---|---|---|---|---|---|---|---|---|
| | Evaluation Agreement | | | | | Evaluation Agreement | | | |
| | Emotion Agreement | 12 | 0.517 | .085 | | Emotion Agreement | 12 | -0.130 | .688 |
| | Group Emotion Agreement | 12 | 0.538 | .071 | | Group Emotion Agreement | 12 | 0.002 | .996 |
| | Individual Performance Evaluation Agreement | 12 | 0.046 | .886 | | Individual Performance Evaluation Agreement | 12 | -0.264 | .408 |
| PC23 | Group Mean | 12 | 0.252 | .430 | PC31 | Group Mean | 12 | -0.189 | .557 |
| | Group Emotion Mean | 12 | 0.366 | .242 | | Group Emotion Mean | 12 | -0.112 | .728 |
| | Emotion Mean | 12 | 0.329 | .296 | | Emotion Mean | 12 | 0.022 | .946 |
| | Individual Performance Evaluation Mean | 12 | 0.366 | .242 | | Individual Performance Evaluation Mean | 12 | -0.139 | .666 |
| | Group Performance Evaluation Agreement | 12 | 0.361 | .249 | | Group Performance Evaluation Agreement | 12 | -0.305 | .335 |
| | Emotion Agreement | 12 | 0.124 | .702 | | Emotion Agreement | 12 | -0.357 | .254 |
| | Group Emotion Agreement | 12 | 0.468 | .125 | | Group Emotion Agreement | 12 | -0.362 | .248 |
| | Individual Performance Evaluation Agreement | 12 | 0.525 | .079 | | Individual Performance Evaluation Agreement | 12 | -0.132 | .683 |
